# mTORC1 promotes P-body assembly to support selective translation of CDS-m^6^A mRNAs

**DOI:** 10.64898/2026.11.26.769392

**Authors:** Hirokazu Nakatsumi, Kazuya Ichihara, Taisuke Mizoo, Atsushi Hatano, Koichi Ogami, Yuichiro Hara, Toru Aonuma, Yuzuki Imai, Akinobu Matsumoto, Masaki Matsumoto, Keiichi I. Nakayama, Michiko Shirane

**Affiliations:** Department of Molecular Biology, Graduate School of Pharmaceutical Sciences, Nagoya City University, Nagoya, Aichi 467-8603, Japan; Division of Biological Science, Graduate School of Science, Nagoya University, Nagoya, Aichi 464-8602, Japan; Department of Biomaterials, Graduate School of Medicine, Dentistry and Pharmaceutical Sciences, Okayama University, Okayama 700-8558, Japan; Department of Omics and Systems Biology, Niigata University, Graduate School of Medicine Dentistry and Health Science, Niigata, Niigata 951-8510, Japan; Division of Molecular Oncology, Center for Neurological Diseases and Cancer, Nagoya University Graduate School of Medicine, Nagoya, Aichi 466-8550, Japan; Research Center for Genome and Medical Sciences, Tokyo Metropolitan Institute of Medical Science, Tokyo 156-8506, Japan; Division of Cell Biology, Medical Institute of Bioregulation, Kyushu University, Fukuoka, Fukuoka 812-8582, Japan; Division of Biochemistry and Integrative Cancer Research, The Institute of Medical Science, The University of Tokyo, Tokyo 108-8639, Japan; Department of Data Science, School of Frontier Engineering, Kitasato University, Sagamihara, Kanagawa 252-0373, Japan; Anticancer Strategies Laboratory, Advanced Research Initiative, Institute of Science Tokyo, Tokyo 113-8510, Japan

## Abstract

Growth and nutrient signals regulate both global protein synthesis and selective mRNA translation through mTORC1. Processing bodies (P-bodies) are membraneless cytoplasmic compartments generally regarded as sites of translational repression and mRNA decay, and whether they contribute to this selectivity is unknown. Here, we show that mTORC1 rapidly promotes P-body assembly and enhances translation of P-body-associated mRNAs enriched for N^6^-methyladenosine within their coding sequences (CDS-m^6^A). mTORC1-stimulated translation of these mRNAs is selectively attenuated in P-body-deficient cells. The m^6^A reader YTHDF2, previously implicated in mRNA decay, localizes to a subset of mTORC1-responsive P-bodies and promotes translation of these transcripts. Unlike canonical 5′TOP mRNAs, which encode the translation machinery, this mRNA population is enriched for transcripts encoding transcriptional regulators and early stimulus-response proteins. These findings identify P-bodies as positive mediators of selective translation downstream of mTORC1 and suggest a route by which growth signals tune the production of regulatory proteins.

## Introduction

Cells continuously adjust their proteome to match their environment. This adjustment occurs in part at the level of translation, which allows rapid and selective changes in protein output without new transcription^1^. The mechanistic target of rapamycin complex 1 (mTORC1) integrates growth factor and nutrient signals to regulate the translation machinery^2^. By phosphorylating the eIF4E-binding proteins (4E-BPs), mTORC1 releases eIF4E to assemble eIF4F, the complex that recruits ribosomes to the mRNA cap^1,2^. Translation of 5′ terminal oligopyrimidine (5′TOP) mRNAs, which encode ribosomal proteins and translation factors, is especially sensitive to this control^3,4^, and is further shaped by the RNA-binding protein LARP1^5,6^. These studies define how mTORC1 acts through initiation factors and RNA-binding proteins.

Processing bodies (P-bodies) are one kind of cytoplasmic RNA granule: membraneless condensates that concentrate RNA and protein, enriched in untranslated mRNAs and in proteins involved in translational repression and mRNA decay^7^. Although initially regarded as sites of mRNA degradation, P-bodies store translationally repressed mRNAs, some of which can return to translation when conditions change^8,9^. The mRNAs retained in P-bodies vary across cell types and cellular states^10^, suggesting that P-bodies could contribute to selective gene expression. P-bodies have nonetheless been associated with translational repression and storage rather than with the promotion of translation. m^6^A is a prevalent internal mRNA modification that influences mRNA fate at multiple levels^11^. Studies of m^6^A within coding sequences (CDS-m^6^A) have connected this modification to P-bodies: CDS-m^6^A induces ribosome pausing and triggers translation-dependent mRNA decay. The target mRNAs are enriched in P-bodies, and the m^6^A reader YTHDF2 contributes to their decay^12^. Similarly, m^6^A-induced ribosome stalling and collisions initiate mRNA degradation^13^. P-body localization of CDS-m^6^A mRNAs is therefore read as a step toward their decay.

YTHDF2 is not confined to this outcome, however: first characterized as a reader that destabilizes m^6^A-modified mRNAs^14^, it can also enhance their translation in cancer cells, through interactions with translation initiation factors or within co-condensates with other proteins^15,16^. Whether growth signals can instead direct these mRNAs into translation — and whether P-bodies serve that end — has not been tested.

Here, we show that mTORC1 rapidly promotes P-body assembly in response to insulin. Ribosome profiling of cells with constitutive mTORC1 activity revealed that mTORC1 enhances translation of P-body-associated CDS-m^6^A mRNAs, and that this effect is selectively attenuated in P-body-deficient cells. YTHDF2 localizes to a subset of insulin-induced P-bodies and supports translation of these transcripts, and those most strongly bound by YTHDF2 show the largest mTORC1-dependent changes in translation. Unlike 5′TOP mRNAs, this population is enriched for transcripts encoding transcriptional regulators and early stimulus-response proteins. Our findings reveal a positive role for P-bodies in selective translation downstream of mTORC1 and suggest a route by which growth signals shape the production of regulatory proteins.

## Results

### mTORC1 promotes P-body assembly

We first asked whether growth-stimulated mTORC1 activation influences P-body assembly. In serum-starved HeLa cells, insulin stimulation for 1 h increased the number of P-bodies, visualized by immunofluorescence for the decapping-complex subunit DCP1A, a standard P-body marker, and pretreatment with the ATP-competitive mTOR kinase inhibitor Torin1 prevented this increase. The per-cell distribution shifted accordingly, with insulin raising the proportion of cells containing numerous P-bodies (Figures 1A, 1D, and 1E). Acute activation of mTOR signaling by a growth stimulus is therefore accompanied by increased P-body assembly.

**Figure 1.**
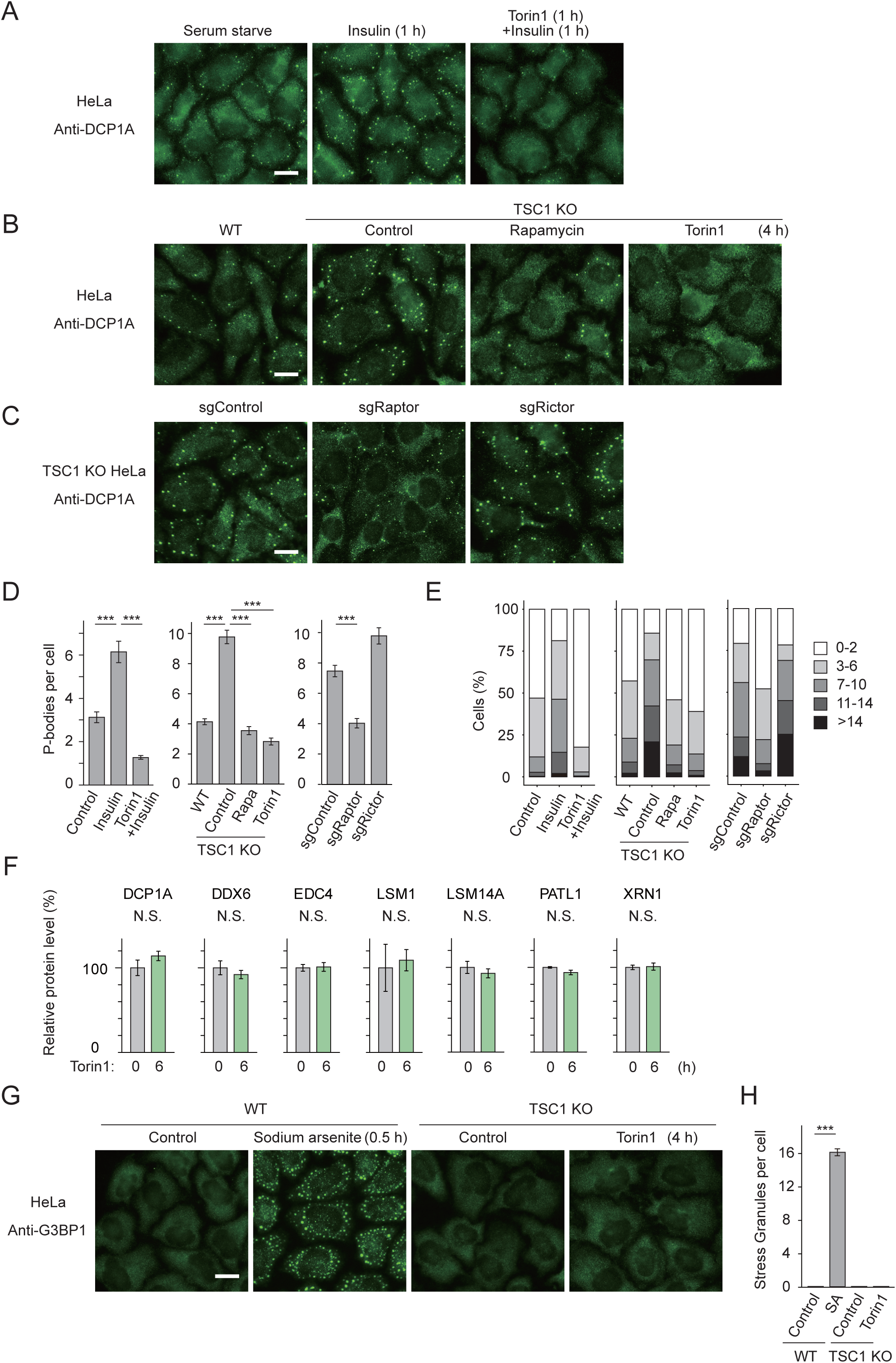
mTORC1 promotes P-body assembly. (A) DCP1A immunofluorescence in serum-starved HeLa cells stimulated with insulin (100 nM, 1 h), with or without Torin1 pretreatment (100 nM, 1 h). (B) DCP1A immunofluorescence in wild-type (WT) and TSC1-knockout (KO) HeLa cells treated with vehicle, rapamycin (10 nM), or Torin1 (100 nM) for 4 h. (C) DCP1A immunofluorescence in TSC1 KO HeLa cells transduced with sgControl, sgRaptor, or sgRictor. (D) DCP1A-positive puncta per cell in (A–C), quantified with CellProfiler^17^. Data are from representative experiments. Bars represent means ± SD across individual cells, with the number of analyzed cells (n) indicated for each condition. Rapa, rapamycin.(E) Percentages of cells containing 0–2, 3–6, 7–10, 11–14, or >14 puncta in (A–C). (F) Protein abundances of DCP1A, DDX6, EDC4, LSM1, LSM14A, PATL1, and XRN1 in TSC1 KO HeLa cells treated with Torin1 (100 nM, 6 h), by quantitative proteomics. Bars, means ± SD normalized to untreated cells (100%); n = 4 biological replicates; two-sided unpaired t tests per protein. (G) G3BP1 immunofluorescence in WT HeLa cells with or without sodium arsenite (500 µM, 30 min), and in TSC1 KO HeLa cells with or without Torin1 (100 nM, 4 h). (H) G3BP1-positive puncta per cell in (G). Bars represent means ± SD across individual cells from representative experiments. Scale bars, 20 µm (A–C and G). ***P < 0.001 in (D and H); N.S., P ≥ 0.05 in (F). See also Figure S1.

To work from a constitutively high baseline rather than a transient stimulus, we next turned to TSC1-knockout (KO) HeLa cells, in which loss of the mTORC1 inhibitor TSC1 leaves mTORC1 signaling constitutively active (Figure S1A). These cells contained more P-bodies than wild-type cells, and either rapamycin or Torin1 reduced their number (Figures 1B, 1D, and 1E). Because Torin1 inhibits both mTORC1 and mTORC2^2,4^, we disrupted the defining subunit of each complex. Disruption of Raptor, which is specific to mTORC1, likewise reduced P-body numbers in TSC1 KO cells, whereas disruption of Rictor, which is specific to mTORC2, did not (Figures 1C–1E and S1A). The agreement between pharmacological and genetic inhibition, together with the selectivity for Raptor over Rictor, indicates that P-body assembly depends on mTORC1 rather than mTORC2.

P-body loss after mTORC1 inhibition could reflect reduced abundance of P-body components or secondary effects of established mTORC1 outputs. Torin1 treatment for 6 h left the total abundance of DCP1A, DDX6, EDC4, LSM1, LSM14A, PATL1 and XRN1 — core P-body constituents spanning the decapping complex, the 5′–3′ decay pathway and factors required for P-body assembly — essentially unchanged (Figure 1F). Two prominent outputs of acute mTORC1 inhibition could account for this loss: LARP1-mediated repression of TOP mRNA translation^5,6^, and the induction of autophagy, which requires ATG13^18^. sgRNAs targeting LARP1 or ATG13 did not prevent Torin1 from reducing DCP1A-positive P-bodies in TSC1 KO cells (Figure S1B). P-body loss is therefore not simply a consequence of depleted P-body components, LARP1-mediated translational control, or ATG13-dependent autophagy.

Finally, we asked whether mTORC1 also controls the assembly of other RNA granules. G3BP1-positive stress granules, a distinct type of cytoplasmic condensate, were readily induced by sodium arsenite, but were neither increased in TSC1-deficient cells nor affected by Torin1 (Figures 1G and 1H).

### mTORC1-associated phosphorylation is enriched among P-body-associated factors

Because P-body loss upon mTORC1 inhibition was not explained by reduced abundance of P-body proteins, we considered post-translational regulation. In condensate components that contain intrinsically disordered regions (IDRs), phosphorylation can govern phase separation and molecular partitioning^19^. Reanalysis of published phosphoproteomic data^20^ showed that serum stimulation increased phosphorylation of DCP1A at T225/S226 and of PATL1 at T51. Rapamycin, but not the MEK inhibitor U0126, suppressed both responses, placing them downstream of mTORC1 rather than the ERK pathway. Insulin stimulation and amino acid readdition also rapidly elicited these responses (Figure 2A). Phosphorylation of DCP1A and PATL1 therefore tracks with mTORC1 activity across several growth and nutrient signals.

**Figure 2.**
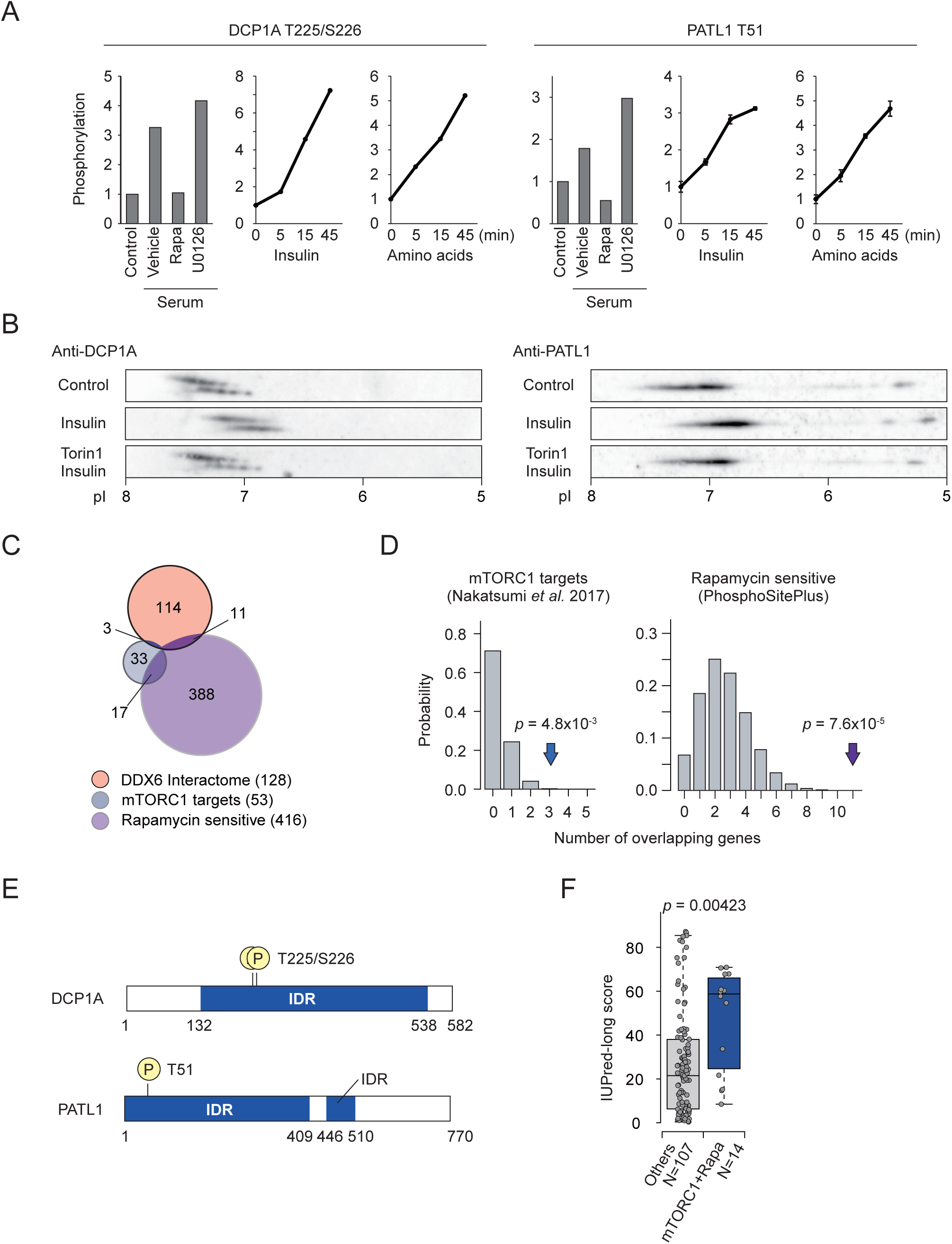
mTORC1-associated phosphorylation is enriched among P-body-associated factors. (A) Phosphorylation of DCP1A T225/S226 and PATL1 T51 from published phosphoproteomics^20^: serum stimulation (20 min) with or without rapamycin (10 nM) or U0126 (10 µM) pretreatment (30 min), and 0, 5, 15, and 45 min after insulin stimulation or amino acid readdition. Data reproduced from the original study; error bars, SD for phosphosites quantified in three or more measurements. (B) Two-dimensional gel electrophoresis and immunoblotting of DCP1A and PATL1 in serum-starved HeLa cells stimulated with insulin (100 nM, 30 min), with or without Torin1 pretreatment (100 nM, 30 min). (C) Overlap among the DDX6 interactome (128 proteins)^21^, mTORC1-associated phosphoproteins (53)^20^, and rapamycin-sensitive phosphoproteins (416; PhosphoSitePlus)^22^. (D) Observed overlaps of 3 and 11 against hypergeometric null distributions. (E) Intrinsically disordered regions (IDRs) and the indicated phosphorylation sites in DCP1A and PATL1. (F) IUPred-long disorder scores^23^ of the 14 DDX6 interactors in either phosphorylation set in (C) versus the remaining 114 (107 with available scores; seven without an annotation were excluded). Each point, one protein; center lines, medians; boxes, 25th–75th percentiles; whiskers, the most extreme values within 1.5 × IQR. The two groups were compared by a two-sided Wilcoxon rank-sum test. See also Table S1.

We assessed these responses independently by two-dimensional electrophoresis. In serum-starved HeLa cells, insulin stimulation for 30 min shifted the spot patterns of both proteins toward lower isoelectric points, as expected for the addition of negatively charged phosphate groups, and Torin1 pretreatment suppressed these shifts (Figure 2B), consistent with the phosphoproteomic data.

The DCP1A and PATL1 responses raised the question of whether mTORC1-associated phosphorylation extends to other P-body-associated factors. As a proxy for the P-body protein network, we used the DDX6 interactome, because DDX6 is a core scaffold around which P-body components assemble^21^. Comparing its 128 members with 53 reported mTORC1-associated phosphoproteins and 416 rapamycin-sensitive phosphoproteins in PhosphoSitePlus^22^, we found three DDX6 interactors in the former set and eleven in the latter (Figure 2C). Both overlaps exceeded chance expectation (Figure 2D), indicating that mTORC1-linked phosphorylation is enriched among DDX6-associated proteins.

These factors also shared a structural feature. DCP1A T225/S226 and PATL1 T51 both lie within IDRs (Figure 2E), and the 14 DDX6 interactors present in either phosphorylation set had higher disorder scores than the remaining 107 with available scores (Figure 2F). mTORC1-associated phosphorylation is thus enriched among P-body-associated proteins with pronounced intrinsic disorder, a context in which phosphorylation can regulate condensate assembly^19^.

### P-body-associated CDS-m^6^A mRNAs exhibit P-body-dependent translational responses

Because P-bodies concentrate a defined subset of mRNAs^9^ and have been linked to their translational repression and decay^7^, mTORC1-dependent changes in P-body assembly could influence the abundance or translation of those transcripts. We therefore sought to identify mRNAs regulated by the mTORC1–P-body axis. A connection between P-bodies and m⁶A offered a starting point: mRNAs carrying CDS-m⁶A can accumulate in P-bodies during translation-dependent decay^12^. We integrated two published measurements: P-body localization scores, the ratio of RNA quantified in P-bodies purified by fluorescence-activated particle sorting to that quantified in the pre-sorted fraction^9^, and m⁶A enrichment scores from m⁶A-seq^24^, for the 5′ untranslated region (5′UTR), coding sequence (CDS), and 3′ untranslated region (3′UTR). Consistent with a previous analysis based on an independent m⁶A map^12^, CDS-m⁶A enrichment, but not m⁶A in either UTR, separated P-body-enriched mRNAs into two populations. We designated the CDS-m⁶A-low population Gate #1 (PB-m⁶A^low^) and the CDS-m⁶A-high population Gate #2 (PB-m⁶A^high^); the remaining mRNAs were assigned to Others (Figure 3A). mTORC1 promotes the translation of TOP mRNAs^3,4^, yet known TOP mRNAs were rare in Gate #1 and absent from Gate #2 (Figure 3B). Gate #1 and Gate #2 are therefore largely distinct from the canonical TOP-mRNA class regulated by mTORC1, and we asked whether either population is regulated by mTORC1 through P-bodies.

**Figure 3.**
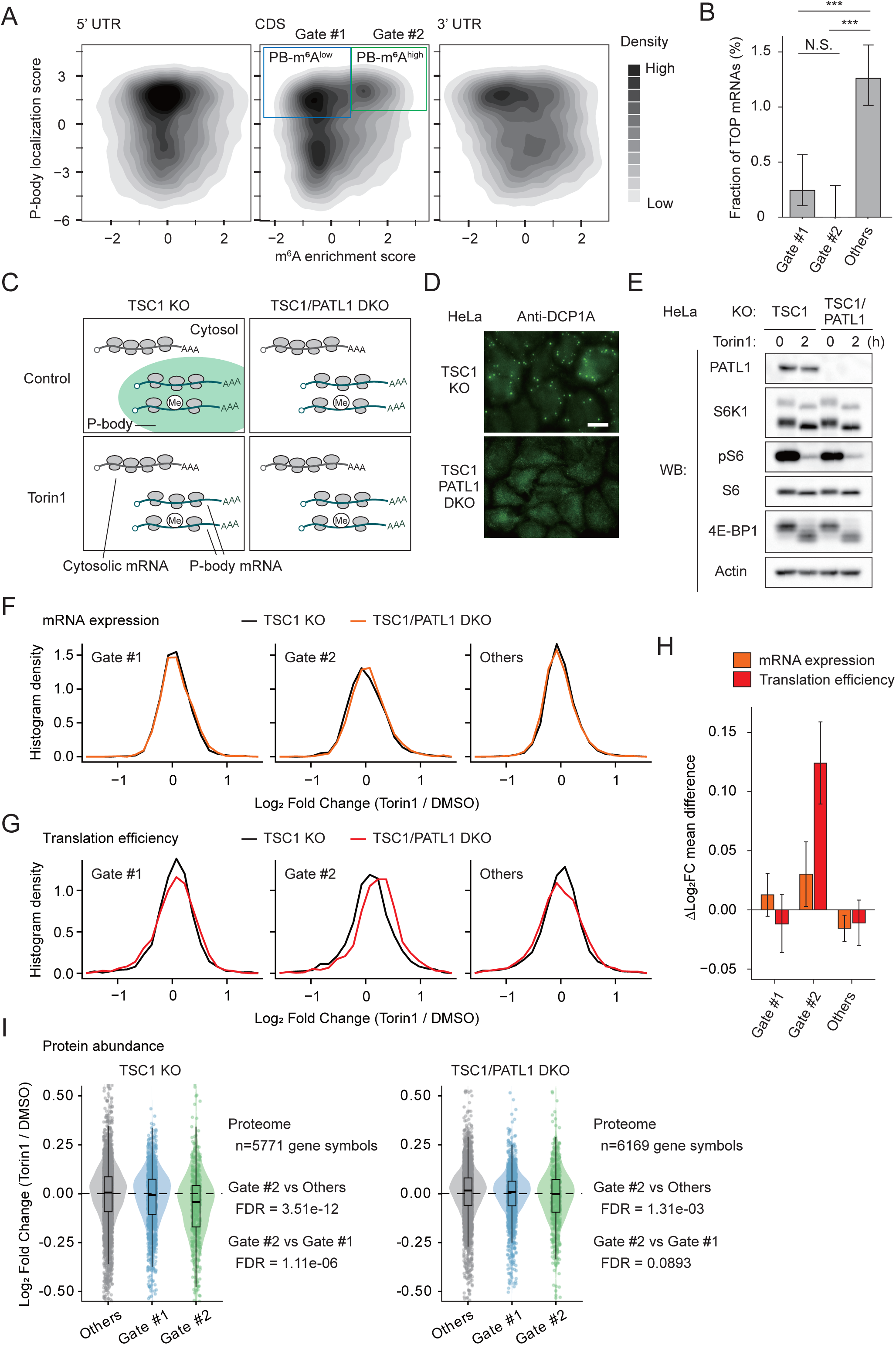
P-body-associated CDS-m^6^A mRNAs exhibit P-body-dependent translational responses. (A) Joint distributions of P-body localization scores^9^ and m^6^A enrichment scores in the 5′UTR, CDS, or 3′UTR^24^. P-body-enriched genes with low or high CDS-m^6^A scores were assigned to Gate #1 or Gate #2, and the remaining genes to Others (11,057 genes: 2,200, 1,548, and 7,309). (B) Proportions of known TOP mRNAs^27^ in each group (86 TOP genes within the 9,821-gene universe). Error bars, Wilson score 95% binomial CIs. Three pairwise two-sided Fisher’s exact tests, Benjamini–Hochberg adjusted; ***adjusted P < 0.001; N.S., adjusted P ≥ 0.05. (C) Schematic of RNA-seq and ribosome profiling in TSC1 KO and TSC1/PATL1 double-knockout (DKO) HeLa cells treated with Torin1 (100 nM, 2 h). The mRNA localization shown is a proposed model. (D) DCP1A immunofluorescence in TSC1 KO and TSC1/PATL1 DKO HeLa cells. Scale bar, 20 µm. (E) Immunoblot analysis of PATL1, S6K1, phosphorylated S6 (pS6), S6, 4E-BP1, and actin under the conditions in (C). (F and G) Distributions of Torin1-induced log2 fold changes in RNA abundance (F) and translation efficiency (TE; G, from the DESeq2 treatment-by-assay interaction coefficient^28^), in the 9,821-gene filtered population (2,061 Gate #1, 1,333 Gate #2, 6,427 Others). TSC1 KO, black; DKO, orange (F) or red (G). (H) Differences in mean log2 fold change (DKO minus TSC1 KO) within each group, for RNA abundance (orange) and TE (red). Error bars, percentile 95% bootstrap CIs. (I) Gene-level protein-abundance log2 fold changes after Torin1 (100 nM, 6 h); n = 5,771 (TSC1 KO) and 6,169 (DKO) gene symbols. Center lines, medians; boxes, IQR; whiskers, observations within 1.5 × IQR. Three two-sided Wilcoxon rank-sum comparisons per genotype, Benjamini–Hochberg adjusted; the FDRs for Gate #2 versus Others and Gate #2 versus Gate #1 are shown. See also Table S2.

To address this, we compared the response to mTORC1 inhibition in cells with and without P-bodies. TSC1-deficient HeLa cells, in which mTORC1 is constitutively active, served as P-body-proficient cells. On this background we generated TSC1/PATL1 double-knockout (DKO) cells to remove P-bodies, since PATL1 is required for P-body assembly^25^ (Figure 3C). DCP1A-positive P-bodies were present in TSC1 KO cells but undetectable in DKO cells, whereas mTORC1 activity and its inhibition by Torin1 were preserved in DKO cells (Figures 3D and 3E). Each genotype was treated with DMSO or Torin1, and mRNA abundance and translation efficiency were measured by RNA-seq and ribosome profiling. Ribosome profiling quantifies ribosome-protected mRNA fragments, and comparison with RNA-seq data allows changes in translation efficiency to be estimated^26^. We analyzed cells after 2 h of Torin1 to capture the early consequences of the change in P-body assembly.

We first examined mRNA abundance. Torin1-induced changes in mRNA abundance were nearly superimposable between the two genotypes for Gate #1, Gate #2, and Others (Figure 3F). The presence of P-bodies therefore had little effect on the response of mRNA abundance to Torin1. We then examined translation efficiency. For Gate #1 and Others, the distributions of Torin1-induced changes overlapped between genotypes, whereas for Gate #2 the distribution in DKO cells was shifted toward higher values relative to TSC1 KO cells (Figure 3G). The translation-efficiency response of Gate #2 thus changes selectively with the presence of P-bodies. The mean genotype-dependent difference in the Torin1 response (DKO minus TSC1 KO) was greater for Gate #2 than for Gate #1 or Others in translation efficiency, but not in mRNA abundance (Figure 3H). Regulation of Gate #2 by the mTORC1–P-body axis therefore appears in translation efficiency rather than in mRNA abundance.

If P-bodies acted solely to repress translation, their reduction upon mTORC1 inhibition would be expected to relieve translational repression of Gate #2 mRNAs in TSC1 KO cells. We therefore measured protein abundance after 6 h of Torin1. Contrary to this prediction, proteins encoded by Gate #2 mRNAs decreased more than those encoded by Gate #1 or Others in TSC1 KO cells, and this preferential decrease was attenuated in P-body-deficient DKO cells (Figure 3I). The translation-efficiency data were consistent with this finding: the Torin1-induced change for Gate #2 was more negative in TSC1 KO cells than in DKO cells, whereas relief of translational repression would predict the opposite (Figure 3G). Together, these results support a model in which mTORC1-dependent P-body assembly sustains the translation of Gate #2 mRNAs, at least in the short term.

### CDS-DRACH motifs and P-bodies shape the mTORC1 response of a reporter

To test whether the CDS-DRACH motifs contribute to this response, we built mCherry reporters with a C-terminal PEST sequence to shorten the protein’s half-life and make steady-state levels more responsive to changes in synthesis. The D(+) reporter retained 13 m^6^A consensus (DRACH, where D = A/G/U, R = A/G, and H = A/C/U) motifs within the CDS, whereas D(−) carried synonymous substitutions disrupting all 13 (Figure 4A). Because the two reporters encode the same protein, this design lets us examine the effect of synonymous substitutions that disrupt the CDS-DRACH motifs.

**Figure 4.**
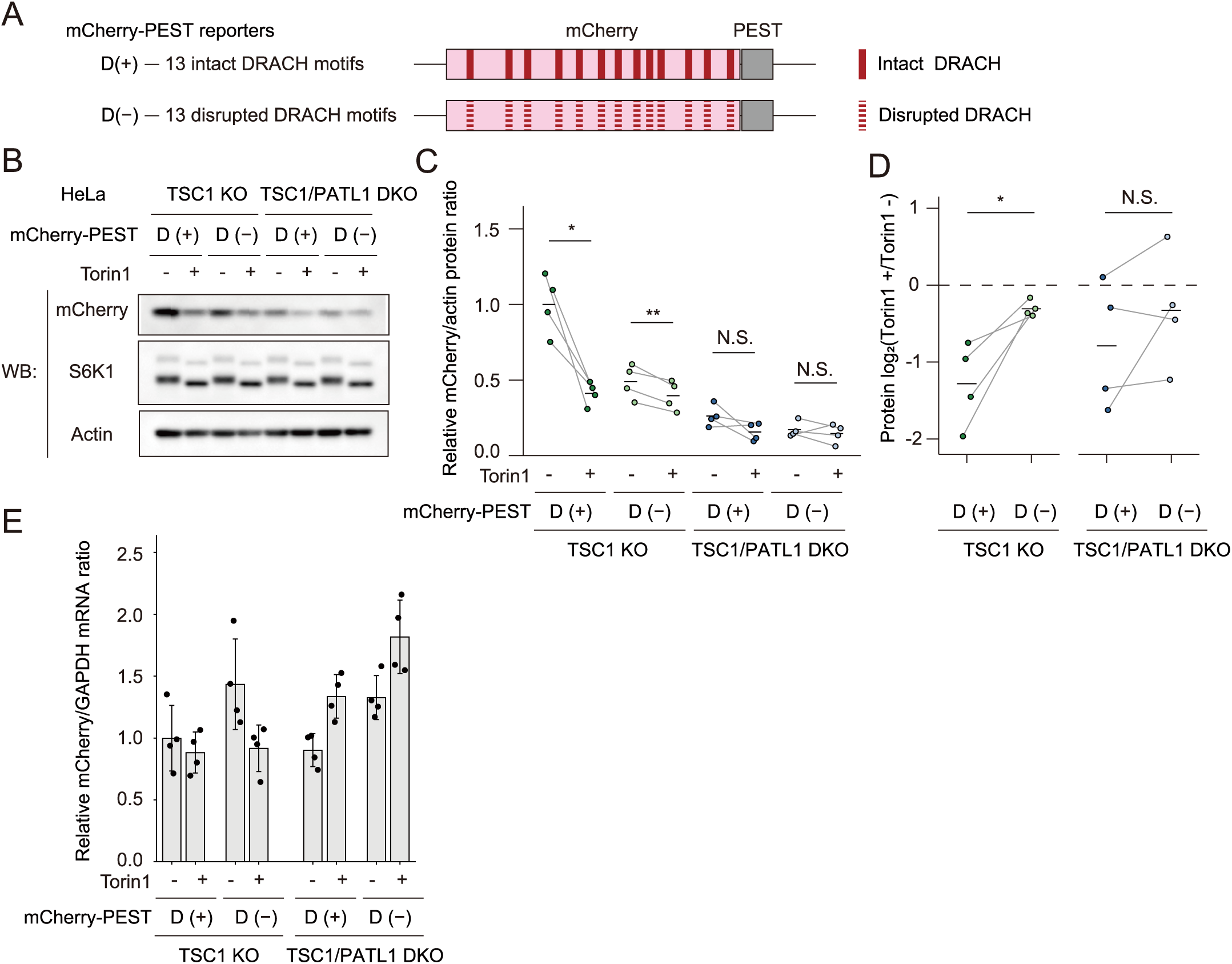
CDS-DRACH motifs and P-bodies shape the mTORC1 response of a reporter. (A) Schematics of the mCherry-PEST D(+) and D(−) reporters. D(+) retains 13 DRACH motifs in the CDS; D(−) carries synonymous substitutions disrupting all 13. Both encode the same amino acid sequence. (B) Immunoblotting of mCherry, S6K1, and actin in TSC1 KO and TSC1/PATL1 DKO HeLa cells expressing D(+) or D(−), with or without Torin1 (100 nM, 12 h). (C) Paired quantification of mCherry/actin ratios, normalized with untreated TSC1 KO D(+) set to 1. Each point, one independent experiment (n = 4) in which all eight conditions were measured; gray lines connect Torin1-treated and untreated samples within an experiment; black bars, means. No SD or SEM is shown. (D) Log2 Torin1 responses, log2[Torin1(+)/Torin1(−)], within each experiment, genotype, and DRACH condition (n = 4). Lines connect D(+) and D(−) within an experiment; black bars, means; dashed line, zero. (E) Reporter mRNA abundance by RT-qPCR, normalized to GAPDH, under the conditions in (B). One experiment with four wells per condition; bars, mean ± SD; no statistical comparisons were made. Two-sided paired t tests compared Torin1-treated versus untreated samples in (C) and D(+) versus D(−) in (D). *P < 0.05; **P < 0.01; N.S., P ≥ 0.05. Untreated reporter abundance between genotypes, not indicated in the figure: P = 0.0112 [D(+)] and 0.0252 [D(−)], unadjusted.

In P-body-proficient (TSC1 KO) cells, the Torin1-induced reduction in mCherry abundance was larger for D(+) than for D(−) (Figures 4B–4D), showing that the CDS-DRACH motifs shape this response. In P-body-deficient (DKO) cells, reporter abundance was low even while mTORC1 was active, for both reporters, and neither reporter showed a detectable response to Torin1 (Figures 4B and 4C). Reporter mRNA abundance did not track these protein differences (Figure 4E), and because D(+) and D(−) encode the same protein, the encoded sequence cannot account for them. These results place DRACH- and PATL1-dependent control of reporter output downstream of transcript abundance, at a step that includes translation.

### YTHDF2 promotes the translation efficiency of P-body-associated CDS-m^6^A mRNAs

Because CDS-DRACH motifs contributed to the reporter response, we next asked whether an m⁶A reader connects the modification to this response. We focused on YTHDF2, which localizes to P-bodies^14^ and has been implicated both in mRNA decay and, in some settings, in translation^15,16^. Immunofluorescence showed that YTHDF2 was enriched in only a subset of DCP1A-positive P-bodies after insulin stimulation (Figure 5A). We therefore classified DCP1A-high puncta into YTHDF2-low and YTHDF2-high groups based on their YTHDF2 signals (Figure 5B). Across three independent experiments, insulin increased the fraction of YTHDF2-high puncta more strongly than that of YTHDF2-low puncta. Torin1 pretreatment reduced both fractions, with a larger decrease in percentage points for the YTHDF2-high group (Figure 5C). These results indicate that YTHDF2-enriched P-bodies show a greater insulin-induced response.

**Figure 5.**
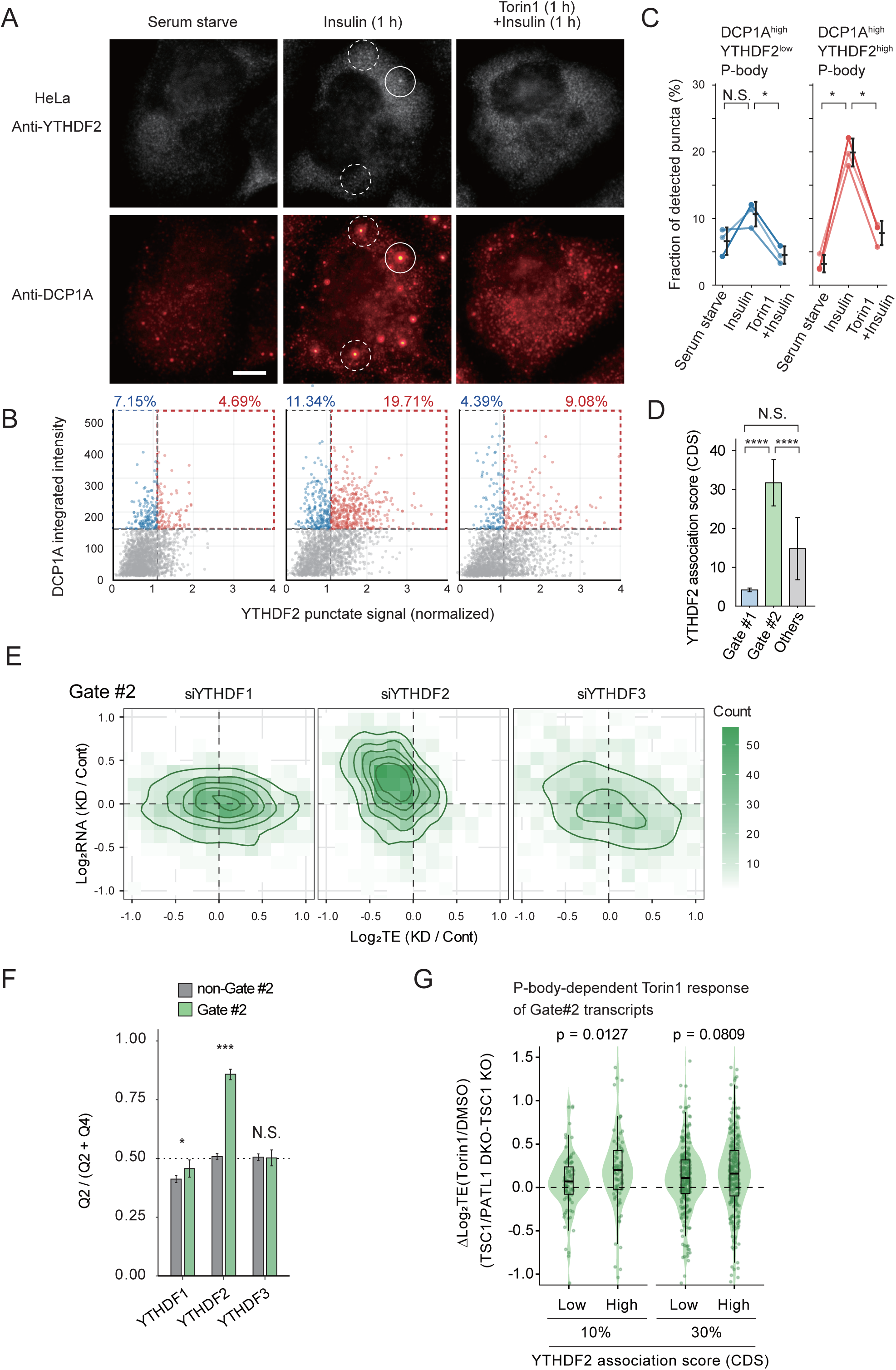
YTHDF2 promotes the translation efficiency of P-body-associated CDS-m^6^A mRNAs. (A) DCP1A and YTHDF2 immunofluorescence in cells treated as in Figure 1A. Solid and dashed circles indicate DCP1A puncta with and without YTHDF2 signal, respectively. Scale bar, 5 µm. (B) DCP1A-high puncta (integrated intensity ≥ 150) classified as YTHDF2-low or YTHDF2-high based on their difference-of-Gaussians signal (threshold ≥ 1.1 after per-experiment normalization). Percentages are of all detected puncta (n = 6,884, 8,609, and 7,092); 2,000 puncta per condition are plotted. One representative experiment of three. (C) Fractions of all detected puncta classified as DCP1A-high/YTHDF2-low (left) or DCP1A-high/YTHDF2-high (right). Each point, one independent experiment (n = 3); lines connect conditions within an experiment; black symbols, mean ± SD. Two-sided paired t tests compared serum starvation with insulin, and insulin with insulin plus Torin1; P values were adjusted across the four comparisons by the Holm method. *Adjusted P < 0.05; ns, not significant. (D) YTHDF2 association scores (Materials and Methods)^14^ for Gate #1 (n = 865), Gate #2 (n = 916), and Others (n = 3,143). Bars, means; error bars, t-based 95% CIs; two-sided Mann–Whitney U tests. ****P < 0.0001; N.S., P ≥ 0.05. (E) Joint distributions of TE and RNA abundance log2 fold changes (knockdown/control) after individual YTHDF1, YTHDF2, or YTHDF3 knockdown^14,29,30^; Gate #2 subsets of 1,356, 1,432, and 1,372 genes. Color, gene counts; contours, density. (F) Q2/(Q2 + Q4) for Gate #2 and non-Gate #2 (Gate #1 plus Others) in (E), where Q2 denotes TE log2FC < 0 with RNA log2FC ≥ 0 and Q4 the converse. Error bars, Clopper–Pearson 95% binomial CIs; two-sided Fisher’s exact tests, unadjusted P values. *P < 0.05; ***P < 0.001; N.S., P ≥ 0.05. (G) Gate #2 genes stratified by YTHDF2 CDS association score; the highest and lowest 10% or 30% (92 and 275 genes per group) were compared for Δlog2TE = log2FC(DKO) − log2FC(TSC1 KO). Points, genes; violins, distributions; boxes, IQR; center lines, medians; whiskers, observations within 1.5 × IQR. Two-sided Wilcoxon rank-sum tests, unadjusted P values. See also Figure S2.

We next asked whether YTHDF2 physically associates with the coding sequences of these mRNAs, using published PAR-CLIP data^14^, which map protein–RNA contacts by UV crosslinking. An abundant transcript can appear strongly bound, so we normalized the CDS signal to input RNA; the resulting association score reflects relative binding per transcript rather than expression level. Association scores were higher in Gate #2 than in Gate #1 or Others (Figure 5D).

To ask whether YTHDF2 supports the translation of these mRNAs, we reanalyzed published ribosome-profiling and RNA-seq data from HeLa cells individually depleted of YTHDF1, YTHDF2, or YTHDF3^14,29,30^; comparing the three paralogs tests whether an effect is specific to YTHDF2. YTHDF2 knockdown lowered translation efficiency and raised mRNA abundance in Gate #2 relative to non-Gate #2, whereas comparable Gate #2-biased effects were not seen for YTHDF1 or YTHDF3 (Figures 5E and 5F). Wang et al. noted a reduced translation efficiency of YTHDF2 targets and attributed it to the accumulation of non-translating mRNA^14^. Because translation can influence mRNA stability, in part through deadenylation-dependent decay^31^, the two responses need not be independent: the rise in mRNA abundance upon YTHDF2 depletion could follow from the fall in translation efficiency. To test whether each response persisted after matching for the other, we drew two non-Gate #2 control sets: one matched to Gate #2 for the YTHDF2-knockdown mRNA-abundance change and the other for the translation-efficiency change. When the control was matched for mRNA-abundance change, Gate #2 still showed a larger fall in translation efficiency; when it was matched for translation-efficiency change, Gate #2 still showed a larger rise in mRNA abundance (Figure S2). Neither change is therefore a simple consequence of the other.

Finally, we tested whether stronger YTHDF2 association was linked to a larger P-body-dependent translational response, as expected if YTHDF2 mediates this effect. Stratifying Gate #2 mRNAs by CDS association score, the between-genotype difference in translation-efficiency response (DKO minus TSC1 KO) was larger for the highest 10% than for the lowest 10%; the 30% cutoff showed the same direction of effect without reaching significance (Figure 5G).

Together, YTHDF2 marks the more mTORC1-responsive P-bodies, associates preferentially with the CDS of these mRNAs, and supports their translation.

### P-body-associated CDS-m^6^A mRNAs are enriched for early stimulus-response genes

Having established a P-body-dependent translational response, we asked what these mRNAs encode, beginning with gene ontology (GO) analysis across the three groups. Gate #2 was markedly enriched for genes involved in RNA polymerase II-dependent transcriptional regulation relative to Gate #1 and Others (Figure 6A). To place this in terms of defined response programs, we performed overrepresentation analysis (ORA) against the 50 Hallmark gene sets^32^, which identified TNFα signaling via NF-κB as the only significant set (Figure 6B). This Hallmark set includes immediate early genes (IEGs) such as FOS and JUN, which are transcribed within minutes of stimulation without new protein synthesis^33^. We therefore asked whether the enrichment reflected a particular transcription-factor program or early-response genes more broadly. To examine these possibilities, we tested 102 transcription-factor target sets^34^, including that of RELA (the p65 subunit of NF-κB), alongside IEGs and delayed primary response genes (DPRGs), which are induced later but, like IEGs, independently of new protein synthesis^33^. Only the IEG set reached significance (Figure 6C). While this does not exclude regulation by individual transcription factors, the data point to early-response genes as a class.

**Figure 6.**
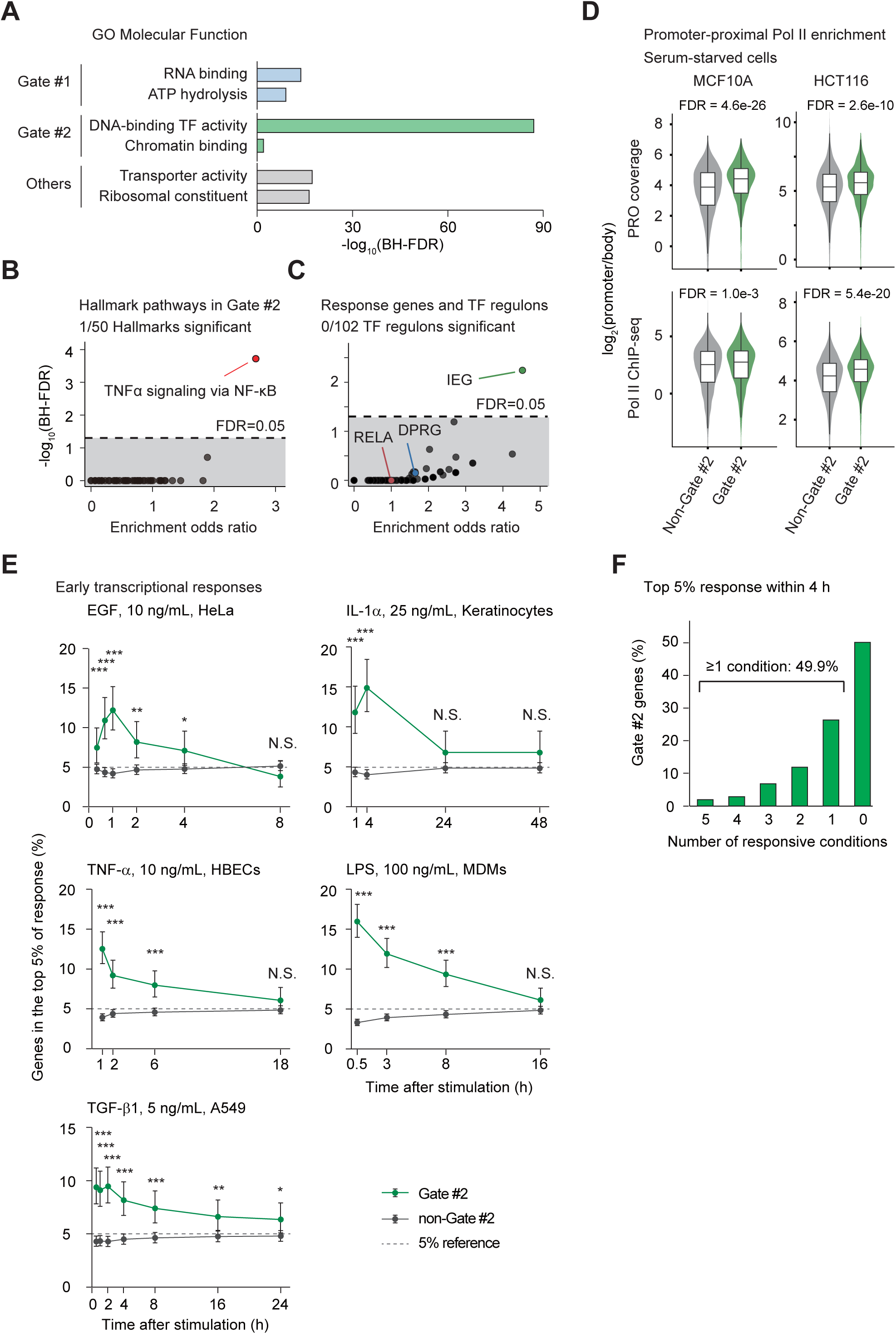
P-body-associated CDS-m^6^A mRNAs are enriched for early stimulus-response genes. (A) GO Molecular Function overrepresentation analysis (ORA) of 1,918 MSigDB 2026.2.Hs gene sets^43–45^ against a fixed 9,821-gene universe, showing two representative terms per group as −log10(BH-FDR). One-sided Fisher’s exact tests, BH-adjusted within each group. (B) ORA of 50 Hallmark gene sets^32^ in Gate #2 (1,333 genes) relative to the remaining genes, as odds ratios and −log10(BH-FDR). One-sided Fisher’s exact tests, BH-adjusted across the 50 sets. (C) ORA of 102 transcription factor regulons^34^, immediate early genes (IEGs), and delayed primary response genes (DPRGs)^33^ in Gate #2, on the axes of (B). IEG, RELA, and DPRG are highlighted. One-sided Fisher’s exact tests within a common BH family of 24,510 tests. (D) Promoter-proximal to gene-body RNA polymerase II signal (log2 ratio) in Gate #2 and non-Gate #2, from public PRO-seq and Pol II ChIP-seq in serum-starved MCF10A and HCT116 cells^36,37^. Violins, gene-level distributions; FDRs are shown. Two-sided Mann–Whitney tests, BH-corrected across five comparisons, including one Pol II ChIP-seq comparison in HCT116 cells that is not displayed. (E) Reanalysis of five public stimulus time courses^38–42^: HeLa/EGF (10 ng/mL), keratinocytes/IL-1α (25 ng/mL), HBECs/TNF-α (10 ng/mL), MDMs/LPS (100 ng/mL), and A549/TGF-β1 (5 ng/mL). Proportions of genes ranking in the top 5% of induction at each time point are compared between Gate #2 and non-Gate #2. Error bars, 95% Wilson CIs; two-sided Fisher’s exact tests with BH adjustment. *Adjusted P < 0.05; **adjusted P < 0.01; ***adjusted P < 0.001; N.S., adjusted P ≥ 0.05. (F) Among 353 Gate #2 genes evaluable in all five datasets, the number of series in which a gene reached the top 5% at any time within 0 < t ≤ 4 h; 176 genes (49.9%) did so in at least one series. In (B) and (C), dashed lines indicate FDR = 0.05; in (E), the nominal 5% threshold.

Rapidly inducible genes often hold RNA polymerase II in a paused state just downstream of the promoter, a configuration termed promoter-proximal pausing that poises immediate inflammatory response genes for activation^35^. Pausing leaves Pol II concentrated near the promoter relative to the gene body. We therefore compared the ratio of promoter-proximal to gene-body Pol II signal between Gate #2 and non-Gate #2, using published PRO-seq, which maps transcriptionally engaged polymerase genome-wide, and Pol II ChIP-seq data from serum-starved MCF10A and HCT116 cells^36,37^. This ratio was higher in Gate #2 (Figure 6D), a promoter-proximal occupancy configuration associated with rapid transcriptional induction.

These steady-state analyses suggested early inducibility but did not measure induction kinetics. We therefore examined five published time courses to test whether Gate #2 genes respond early across different stimuli and cell types: EGF-stimulated HeLa cells, IL-1α-stimulated keratinocytes, TNF-α-stimulated human bronchial epithelial cells (HBECs), LPS-stimulated monocyte-derived macrophages, and TGF-β1-stimulated A549 cells^38–42^. Early after stimulation, the proportion of genes ranking in the top 5% for induction was significantly higher in Gate #2 than in non-Gate #2 in all five conditions, and the difference narrowed at later time points (Figure 6E). This behavior was not confined to a few genes: among the 353 Gate #2 genes evaluable in all five datasets, 176 (49.9%) reached the top 5% in at least one condition within 4 h (Figure 6F).

P-body-associated CDS-m^6^A mRNAs are thus enriched for transcriptional regulators and encompass a broad set of genes that respond early to diverse extracellular stimuli.

## Discussion

mTORC1 promotes P-body assembly, and P-bodies in turn support the translation of mRNAs marked by m^6^A within their coding sequences — a positive role for P-bodies in translational control downstream of a growth signal, alongside their established functions in repression and decay. The affected transcripts are largely distinct from the 5′TOP mRNAs that encode the translation machinery. The separation between 5′TOP and CDS-m^6^A-enriched transcripts suggests that mTORC1 coordinates two outputs through distinct mechanisms: the translation apparatus itself, and the regulatory proteins through which cells respond to their environment.

mTORC1 inhibition reduces P-bodies despite suppressing translation initiation, which generally favors their assembly^7^. This suggests that mTORC1 promotes P-body assembly through a mechanism beyond its effects on translation initiation. Phosphorylation of intrinsically disordered regions (IDRs) can promote condensate assembly by altering their electrostatic and multivalent interactions. For example, phosphorylation of the disordered repeats of Ki-67 enhances phase separation by changing their charge distribution^46^, while phosphorylation of the C-terminal IDR of FMRP promotes RNA-dependent phase separation in vitro^47^. More directly relevant to P-bodies, JNK-mediated phosphorylation of 4E-T promotes the formation of larger P-bodies under oxidative stress^48^. Together with our findings on DCP1A, PATL1, and DDX6-interacting proteins, these observations support a model in which mTORC1 promotes P-body assembly by modulating interactions among P-body components through phosphorylation.

mTORC1 also promotes m⁶A deposition through WTAP and S-adenosylmethionine, thereby influencing mRNA decay and protein synthesis^49,50^. Because m⁶A is deposited predominantly during transcription^51^, this pathway can affect a broad range of newly synthesized mRNAs, including those modified in the CDS and 3′UTR, with regulatory consequences that may extend over longer periods. In contrast, mTORC1-dependent P-body assembly appears to sustain the translation of a more restricted population, PB-m⁶A^high^ mRNAs, during the early response to changes in mTORC1 signaling. These mechanisms thus differ not only in their temporal and transcript-specific effects but also in the nature of their regulation: one shapes the m⁶A modification landscape, whereas the other influences how modified mRNAs are regulated.

Together, they suggest that mTORC1 may shape the fate of m⁶A-modified mRNAs through both their modification state and the P-body environment.

Previous studies have linked CDS-m⁶A to translation-dependent mRNA decay^12,13^, whereas our findings reveal a P-body-dependent translational response during the early phase of mTORC1 inhibition. These observations need not be mutually exclusive, as CDS-m⁶A-dependent decay itself requires translation. We found that YTHDF2 enrichment varies among P-bodies and changes with mTORC1 signaling, suggesting that P-body composition may be dynamically regulated. Such heterogeneity is consistent with the selective association of PCBP2 with a subset of P-bodies reported previously^52^. Such changes in P-body composition could reshape the relationship between translation and decay over the course of a cellular response, potentially reconciling these distinct observations.

Although P-body-associated mRNAs are translationally repressed, their return to translation after leaving P-bodies has been demonstrated in yeast^8,9^. YTHDF2 was originally shown to direct m⁶A-modified mRNAs toward P-body-associated decay^14^. More recently, CRTC2 was reported to recruit YTHDF2 and promote the transfer of m⁶A-modified mRNAs from decay sites to polysomes through PABP1^16^, whereas YTHDF2 was shown to promote translation through interactions with eIF3F and DDX1^15^. Together, these studies suggest that the protein environment of YTHDF2-bound mRNAs can favor either decay or translation. This raises the possibility that mTORC1-dependent P-body assembly facilitates the transition of CDS-m⁶A mRNAs from a nontranslating to a translating state.

The mTORC1–P-body axis may be particularly relevant to rapid stimulus responses, as its target mRNAs are enriched for transcriptional regulators and immediate early genes. This suggests two layers of control acting in series: external stimuli determine which mRNAs are produced, whereas mTORC1 and P-bodies influence how efficiently those mRNAs are translated. The same stimulus could therefore yield different protein outputs depending on the growth and nutrient state of the cell.

### Limitations of the study

The molecular mechanism by which mTORC1 controls P-body assembly remains undefined; our phosphorylation data are correlative and do not establish a direct action of mTORC1 on specific sites. P-body-deficient cells were generated by deleting PATL1, so effects attributed to P-bodies may also arise from P-body-independent functions of PATL1 itself. We also do not resolve how translational use and decay of P-body-associated mRNAs relate to one another, whether in time or in mechanism, for any individual transcript. Finally, although insulin stimulation altered P-body abundance, we did not test whether the mTORC1–P-body axis changes protein output during an endogenous stimulus-response program.

## Materials and Methods

Reagents and resources used in this study are listed in Table S3.

### Cell culture

HeLa cells were maintained in DMEM supplemented with 10% fetal bovine serum (FBS) and penicillin-streptomycin at 37°C in a humidified incubator with 5% CO₂.

### Growth factor stimulation and pharmacological treatments

Torin1 and rapamycin were used at 100 nM and 10 nM, respectively. For serum starvation, cells were cultured in medium containing 0.1% serum for 16 h. Human recombinant insulin was used at 100 nM. For Figure 1, cells were stimulated with insulin for 1 h with or without Torin1 pretreatment for 1 h; TSC1 KO cells were treated with rapamycin or Torin1 for 4 h; and sodium arsenite was used at 500 μM for 30 min in the stress-granule comparison. For Figure 3, Torin1 treatment lasted 2 h for RNA-seq/ribosome profiling and 6 h for proteomics.

### CRISPR-Cas9-mediated perturbation

sgRNAs were cloned into pX330-U6-Chimeric_BB-CBh-hSpCas9 (Addgene plasmid #42230)^53^. Knockout cells were isolated by limiting dilution, and each TSC1 KO and TSC1/PATL1 DKO line used in this study was derived from a single clone. Loss of protein expression was confirmed by immunoblotting.

### Immunofluorescence and P-body quantification

Cells were fixed with 4% paraformaldehyde, permeabilized with 0.5% Triton X-100, and blocked with 3% FBS. P-bodies were stained with an anti-DCP1A antibody (1:1000) and Alexa Fluor 488-or Alexa Fluor 555-conjugated secondary antibodies. Nuclei were stained with DAPI. Images for P-body counting in Figures 1A–1C were acquired using an OLYMPUS microscope with a 40× objective and analyzed using CellProfiler^17^. DAPI and anti-desmin staining defined nuclei and cell boundaries, respectively, and DCP1A-positive puncta were counted per cell. Threshold settings were applied uniformly across conditions. More than 100 cells from random fields were analyzed per condition.

### DCP1A and YTHDF2 puncta classification

For Figures 5A–5C, DCP1A and YTHDF2 immunofluorescence images were acquired using a 60× objective in three independent experiments. DCP1A puncta were identified using CellProfiler^17^ with speckle enhancement and adaptive Otsu thresholding (object diameter, 5–60 px). Puncta with DCP1A integrated intensity ≥150 were classified as DCP1A-high. YTHDF2 signals were quantified using a difference-of-Gaussians (DoG) filter (σ = 1.4 and 4.0 px).

YTHDF2 DoG scores were normalized using experiment-specific scaling factors empirically selected to align YTHDF2-positive fractions across experiments. Classification thresholds were determined separately for each experiment, and the same parameters were applied to all treatment conditions within each experiment. Puncta from all fields and wells were pooled for each condition, and percentages were calculated relative to all detected puncta.

### Immunoblotting and reporter protein quantification

Cells were lysed in Triton lysis buffer. Equal amounts of protein were separated by SDS-PAGE and transferred to PVDF membranes. Membranes were blocked with skim milk and incubated with primary and HRP-conjugated secondary antibodies. Signals were detected using SuperSignal West Pico Plus substrate and visualized using a Fusion SOLO.7S imaging system. Band intensities were quantified using Fusion software. Four independent experiments were performed, each comprising eight combinations of genotype, DRACH status, and Torin1 treatment. mCherry/actin ratios were divided by the mean across the eight conditions within each experiment, then multiplied by a common factor to set the four-experiment mean for untreated TSC1 KO D(+) cells to 1.

For Figure 4C, log2-transformed normalized mCherry/actin ratios were compared between Torin1-treated and untreated samples within each genotype and DRACH condition using two-sided paired t tests, paired by experiment. For Figure 4D, the log2 Torin1 response was calculated as log2[Torin1(+)/Torin1(−)] within each experiment, genotype, and DRACH condition. D(+) and D(−) responses were compared within each genotype using two-sided paired t tests. P values displayed in Figures 4C and 4D were unadjusted.

### Two-dimensional electrophoresis and phosphoproteomics

DCP1A and PATL1 were analyzed by two-dimensional electrophoresis and immunoblotting in serum-starved HeLa cells stimulated with insulin (100 nM, 30 min), with or without Torin1 pretreatment (100 nM, 30 min).

Phosphorylation signals for DCP1A T225/S226 and PATL1 T51 (Figure 2A) were obtained from Nakatsumi et al.^20^. The conditions included serum stimulation for 20 min, pretreatment with rapamycin (10 nM) or U0126 (10 μM) for 30 min, and insulin or amino acid responses at 0, 5, 15, and 45 min.

### Reporter construction and quantitative RT-PCR

mCherry-PEST-DRACH(+)/(−) reporters were generated by inserting mCherry coding sequences fused to a C-terminal PEST sequence into pcDNA3. The mCherry-PEST-DRACH(−) reporter was generated using a synthetic gene containing synonymous mutations in all 13 DRACH motifs. For the experiments in Figure 4, TSC1 KO and TSC1/PATL1 DKO HeLa cells were transiently transfected with the reporter plasmids using X-tremeGENE 9 DNA Transfection Reagent. Torin1 (100 nM) or vehicle was added 12 h after transfection, and cells were harvested 12 h later.

Total RNA was extracted using RNAiso. cDNA was synthesized using ReverTra Ace qPCR RT Master Mix with gDNA Remover. Quantitative PCR was performed using THUNDERBIRD Next SYBR qPCR Mix on a StepOnePlus Real-Time PCR System. Gene expression levels were normalized to GAPDH and calculated using the ΔΔCt method^54^.

### RNA sequencing and ribosome profiling

TSC1 KO and TSC1/PATL1 DKO HeLa cells were treated with DMSO or Torin1 (100 nM) for 2 h, with two biological replicates per condition. Cells were washed with ice-cold PBS and lysed in buffer containing 20 mM Tris-HCl (pH 7.5), 150 mM NaCl, 5 mM MgCl₂, 1 mM DTT, 1% Triton X-100, and 100 μg/mL cycloheximide. After clarification by centrifugation, separate aliquots of the same lysate from each biological replicate were used for RNA sequencing and ribosome profiling.

For RNA sequencing (RNA-seq), total RNA was extracted from the clarified lysate aliquot using the PureLink RNA Mini Kit (Thermo Fisher Scientific), and RNA quality was assessed using a 2100 Bioanalyzer (Agilent). Poly(A) RNA was isolated using the NEBNext Poly(A) mRNA Magnetic Isolation Module (New England Biolabs). Libraries were prepared using the NEBNext Ultra Directional RNA Library Prep Kit for Illumina (New England Biolabs) and sequenced on an Illumina NovaSeq platform.

For ribosome profiling, clarified lysate containing 10 μg of total RNA was digested with RNase I (20 U) at 25°C for 45 min. Digestion was terminated by adding SUPERase•In RNase inhibitor (Invitrogen) at a final concentration of 20 U/mL, and ribosomes were recovered by ultracentrifugation through a 1 M sucrose cushion. RNA was extracted from the ribosomal pellet using ISOGEN-LS (Nippon Gene) and purified using the Direct-zol RNA Microprep Kit (Zymo Research). Ribosome-protected fragments (RPFs) of 17–34 nt were isolated by denaturing polyacrylamide gel electrophoresis and treated with T4 polynucleotide kinase to repair 2′,3′-cyclic phosphate termini. The fragments were then ligated to preadenylated linkers using truncated T4 RNA ligase 2 (K227Q). Ribosomal RNA was depleted using Ribo-Zero Gold (Illumina). The remaining RNA was reverse-transcribed using ProtoScript II reverse transcriptase (New England Biolabs), and the resulting cDNA was circularized using CircLigase II (Lucigen). Libraries were amplified by PCR, purified by gel electrophoresis, and sequenced on an Illumina NovaSeq platform.

Adapter sequences were trimmed from raw RNA-seq and ribosome-profiling reads. Reads matching gel-marker sequences were removed using Bowtie2^55^ with a custom reference, followed by removal of reads mapping to rRNA and tRNA sequences using Bowtie2. Read quality was assessed using FastQC and MultiQC^56^. The remaining reads were aligned to the human reference genome (hg38) using STAR^57^. Gene-level counts were generated using featureCounts v2.0.3^58^ with the hg38.ensGene.gtf annotation, aggregating reads by gene_id. RNA-seq reads were assigned to exons using unstranded counting, whereas RPF reads were assigned to coding sequences using strand-specific counting (-s 1).

Torin1-induced changes in translation efficiency (TE) were estimated using DESeq2 interaction models^28^. RNA-seq and RPF count matrices were matched by sample identifiers, and genes present in both datasets were retained. TSC1 KO and TSC1/PATL1 DKO cells were analyzed separately. Within each genotype, genes with a total count of at least 10 across the combined RNA-seq and RPF libraries were retained. Counts were rounded to integers, and DESeq2 models were fitted using the design ∼ treatment + SeqType + treatment:SeqType, with DMSO and RNA as the reference levels. The treatmentTorin1.SeqTypeRPF interaction coefficient was extracted as the Torin1-induced change in TE, representing the difference between the model-based RPF and RNA treatment log2 fold changes. Default DESeq2 size-factor estimation and Wald tests were used. Biological sample identifiers and batch were not included as covariates, and no log2 fold-change shrinkage was applied.

For Figure 3H, the response difference within each gene set was calculated as mean(TSC1/PATL1 DKO log2FC) − mean(TSC1 KO log2FC), after separately excluding missing estimates. The two vectors were pooled, and group labels were permuted 100,000 times. Unadjusted two-sided empirical P values were calculated as the fraction of permutations with an absolute mean difference at least as large as the observed difference, without a plus-one correction. Each vector was independently resampled 10,000 times with replacement (seed 123) to obtain percentile 95% bootstrap confidence intervals. The resampling unit was the gene-level response estimate.

### Quantitative proteomic analysis by DIA–LC–MS/MS

Cell lysates containing 50 µg of protein were reduced with 2.5 mM tris(2-carboxyethyl) phosphine hydrochloride (Thermo Fisher Scientific) for 30 min at 37°C and alkylated with 5 mM 2-iodoacetamide (Sigma-Aldrich) for 30 min at room temperature in the dark. Proteins were precipitated with acetone, resuspended in 25 mM triethylammonium bicarbonate by sonication, and digested with trypsin (Laurus Bio). The resulting peptides were desalted with SDB-XC StageTips (3M). Peptides (200 ng per sample) were loaded onto a trap column (L-column2 micro, CERI) and separated on an in-house–fabricated 20-cm analytical column packed with 2-µm octadecylsilane particles (CERI) with a Dionex UltiMate 3000 HPLC system (Thermo Fisher Scientific). Elution was performed with a linear gradient of 5% to 35% solvent B over 70 min at a flow rate of 200 nl/min (solvent A, 0.1% formic acid in water; solvent B, 0.1% formic acid in acetonitrile). The column temperature was kept at 42°C with a column oven (AMR), and eluted peptides were ionized with a nano-electrospray source. A Q Exactive Plus hybrid quadrupole-Orbitrap mass spectrometer (Thermo Fisher Scientific) was operated in DIA mode with positive polarity, and all data were acquired in profile mode. MS1 spectra were collected over a mass-to-charge ratio (*m/z*) range of 430 to 860 at a resolution of 35,000, with an automatic gain control (AGC) target of 1 × 10⁶ and a maximum injection time of 50 ms. MS2 spectra were collected from *m/z* 200 at a resolution of 17,500, with an AGC target of 1 × 10⁶ and the maximum injection time set to auto. Twenty-one DIA isolation windows of 20 *m/z* with a 1-*m/z* overlap were set to cover the *m/z* range of 430 to 850. The normalized collision energy was set to 25%.

For analysis on the Orbitrap Exploris 480, the resulting digests were loaded onto Evotip Pure (Evosep) according to the manufacturer’s instructions. Peptides were separated on an analytical column (EV1109, Evosep) with the preprogrammed 60-samples-per-day gradient on an Evosep One system (Evosep). Eluted peptides were ionized through an emitter (Ref. 20-05 LOTUS emitter, AMR) and introduced into an Orbitrap Exploris 480 mass spectrometer (Thermo Fisher Scientific) equipped with a FAIMS Pro interface (high-field asymmetric waveform ion mobility spectrometry; Thermo Fisher Scientific). The mass spectrometer was operated in DIA mode with positive polarity, and all data were acquired in profile mode. The spray voltage was set to 2.2 kV, the FAIMS compensation voltage to −45 V, the RF lens level to 40%, and the ion transfer tube temperature to 275°C. Full MS spectra were acquired over an *m/z* range of 400 to 1200 at a resolution of 120,000 (at *m/z* 200), with a normalized AGC target of 100% and a maximum injection time of 45 ms. MS2 spectra were acquired over an *m/z* range of 145 to 1450 at a resolution of 30,000, with a normalized AGC target of 1000% and the maximum injection time set to auto. Thirty-four DIA isolation windows of 12 *m/z* with a 1-*m/z* overlap were set to cover the *m/z* range of 480 to 885. The normalized collision energy was set to 28%. All DIA raw data were processed with DIA-NN (version 1.8.1)^59^ in library-free mode against the human UniProt sequence database^60^ (downloaded on April 27, 2021).

For Figure 3I, proteins were assigned to Gates using the 11,057-gene classification without applying the RNA/RPF expression filter. Entries required valid gene symbols and Gate annotations, a nonmissing log2 fold change, a positive log10-transformed average of mean protein abundances at 0 and 6 h, and four quantified measurements at each time point. Log2 fold changes were calculated from the ratio of mean abundance at 6 h to that at 0 h. Abundance rows were matched to Gate annotations by gene symbol, retaining the first annotation record for each nonempty symbol. The analysis unit was a unique gene-symbol abundance row, without ENSG aggregation; 5,771 TSC1 KO and 6,169 TSC1/PATL1 DKO entries were included. Two-sided Wilcoxon rank-sum tests used exact=FALSE and the default continuity correction, with Benjamini–Hochberg correction across three comparisons within each genotype^61^.

### DDX6 overlap and disorder analyses

The DDX6-associated protein set was defined from Supplemental Table 1 of Ayache et al.^21^ by retaining proteins identified in RNase-treated DDX6 immunoprecipitates with a Mascot score ≥100, a threshold set in this study. The mTORC1-associated phosphoprotein, rapamycin-sensitive phosphoprotein, and DDX6 interactome lists contained 53, 416, and 128 unique entries, respectively^20–22^. Rapamycin-sensitive phosphoproteins were identified from PhosphoSitePlus annotations retrieved on March 19, 2026^22^. Empty entries and surrounding whitespace were removed, and gene symbols were converted to uppercase and deduplicated. Lists were intersected using exact gene-symbol matches without alias conversion. Overlap enrichment was assessed using upper-tail hypergeometric probabilities with a background of 20,000 genes.

The IUPred-long disorder score^23^, defined as the percentage of residues predicted to be disordered and obtained from MobiDB^62^ (records retrieved on November 14, 2026), was compared between the 14 overlapping DDX6-interacting proteins and the remaining 107 proteins with available scores using a two-sided Wilcoxon rank-sum test. Seven non-overlapping proteins lacked this annotation and were excluded without imputation.

### P-body localization and CDS-m⁶A integration

P-body localization and CDS-m⁶A data were obtained from Hubstenberger et al.^9^ (E-MTAB-5477; HEK293 cells) and Liu et al.^24^ (GSE46705; HeLa cells), respectively. The common Gate classification comprised 11,057 genes: 2,200 in Gate #1, 1,548 in Gate #2, and 7,309 in Others. Gate #2 was defined by P-body log2FC >1.3 and CDS-m⁶A log2FC >0.5; Gate #1 by P-body log2FC >1.0 and CDS-m⁶A log2FC <0.5. Remaining genes were assigned to Others.

After excluding genes with insufficient signal in the P-body-localization, CDS-m⁶A, RNA-seq, or RPF datasets, 9,821 genes (2,061 Gate #1, 1,333 Gate #2, and 6,427 Others) were used for Gate-stratified RNA-seq and ribosome-profiling analyses. Proteomics used the 11,057-gene classification with assay-specific quantification criteria. Table S2 provides the classification and an ENSG-based indicator of membership in the RNA/RPF-filtered population. This inclusion filter was distinct from the count-sum filter used for DESeq2 model fitting.

### TOP mRNA enrichment

Known 5′TOP mRNAs were obtained from Philippe et al.^27^. Alias expansion of the 97 listed genes yielded 119 identifiers, of which 86 were present in the 9,821-gene universe: 5 in Gate #1, 0 in Gate #2, and 81 in Others. TOP-mRNA proportions were compared between each pair of Gate groups using two-sided Fisher exact tests on TOP versus non-TOP counts, with Benjamini–Hochberg correction across the three comparisons. Error bars represent Wilson score 95% binomial confidence intervals.

### YTHDF knockdown data processing

Public RNA-seq and RPF data were analyzed for individual YTHDF knockdowns. YTHDF1 data comprised control RPF SRR1661491 and RNA SRR1661492, and knockdown RPF SRR1661493 and RNA SRR1661494^29^. YTHDF2 data comprised control RNA SRR944657 and RPF SRR944659, and knockdown RNA SRR944660 and RPF SRR944662^14^. YTHDF3 data comprised control RPF SRR5099275 and RNA SRR5099276, knockdown replicate 1 RPF SRR5099277 and RNA SRR5099278, and knockdown replicate 2 RPF SRR5099279 and RNA SRR5099280^30^.

For Figures 5E, 5F, S2A, and S2B, genes from the 11,057-gene Gate set were retained if RNA and RPF CPM >1 in both control and knockdown samples. For YTHDF3, knockdown CPMs were averaged across replicates before filtering. For YTHDF2, featureCounts v2.0.3^58^ assigned reads to exons by gene_id using hg38.ensGene.gtf, with RNA settings -s 1/-Q 10 and unique-mapping RPF settings -s 0/-Q 0. RNA and RPF libraries were normalized separately using edgeR TMM^63,64^. RNA changes were calculated as knockdown-minus-control differences in logCPM using cpm(log=TRUE, prior.count=0.25). For YTHDF1 and YTHDF2, TE was calculated as log2(RPF CPM + 0.5) − log2(RNA CPM + 0.5), and TE changes as knockdown-minus-control differences. For YTHDF3, CPM values from the two knockdown replicates were averaged for expression filtering; linear per-replicate RPF/RNA ratios were averaged before log2 transformation, without the same pseudocount.

The final YTHDF1, YTHDF2, and YTHDF3 subsets contained 1,356/1,432/1,372 Gate #2 genes and 8,432/8,663/8,451 non-Gate #2 genes, respectively.

### YTHDF distributions and quadrant analysis

TE and RNA abundance changes were displayed as gene-level distributions. For Figure S2B, mean differences were calculated as Gate #2 minus non-Gate #2. Genes were independently resampled within each group with replacement 2,000 times (seed 1234); the 2.5th and 97.5th percentiles defined 95% bootstrap confidence intervals.

Q2 was defined by TE log2FC <0 and RNA log2FC ≥0, and Q4 by TE log2FC ≥0 and RNA log2FC <0. Q2/(Q2+Q4) was calculated using only genes in these quadrants. Exact Clopper– Pearson 95% binomial confidence intervals were calculated using binom.test. Gate #2 and non-Gate #2 proportions were compared using two-sided Fisher exact tests with unadjusted P values.

### YTHDF2 CDS association scores

YTHDF2 CDS association scores were calculated from CDS PAR-CLIP TPM and input RNA TPM^14^. Gene-ID version suffixes were removed, the datasets were merged, and missing numeric measurements were replaced with zero. The association score was (CDS PAR-CLIP TPM + 0.01)/(input RNA TPM + 0.01). Genes were retained if the total mutation count across CDS-assigned PAR-CLIP peaks was >0, input TPM >1, and the score was finite, yielding 865 Gate #1, 916 Gate #2, and 3,143 Others genes. Means and t-distribution 95% confidence intervals were calculated across genes.

For Figure 5G, the highest and lowest 10% or 30% of the 916 eligible Gate #2 genes were selected by YTHDF2 CDS association score (92 or 275 genes per group, respectively). The Torin1 TE response difference was defined as TSC1/PATL1 DKO minus TSC1 KO. Groups were compared using two-sided Wilcoxon rank-sum tests (exact=FALSE), with unadjusted P values.

### Bidirectional matching of RNA and TE changes

The matching analysis included 1,280 Gate #2 genes (PB-m⁶Ahigh) and 8,044 candidate non-Gate #2 genes (1,997 PB-m⁶Alow and 6,047 Others). Genes were required to have nonmissing basal RNA abundance, log2 CDS length, RNA change, and TE change.

CDS length was defined as the largest summed CDS length among annotated transcripts of each gene in hg38.ensGene.gtf and was log2-transformed. Both matching directions used control RNA logCPM and log2 CDS length, together with RNA change for RNA matching or TE change for TE matching. Covariates were centered and scaled to unit sample standard deviation over the combined target and control pool, and Euclidean distances were calculated in the resulting three-dimensional space. Targets were ordered by decreasing distance to their nearest control in the full candidate pool. Each target was assigned the nearest unused control by greedy 1:1 matching without replacement or a caliper. Each direction yielded 1,280 matched gene pairs.

TE changes after RNA matching and RNA changes after TE matching were compared using two-sided paired Wilcoxon signed-rank tests on matched gene pairs (exact=FALSE, with the default continuity correction). P values were unadjusted. Gene-level distributions were visualized as kernel density estimates using ggplot2.

### GO Molecular Function and gene set overrepresentation

Figure 6 overrepresentation analyses used a fixed universe of 9,821 genes, comprising 1,333 Gate #2 and 8,488 non-Gate #2 genes. Gene symbols were mapped using the HGNC snapshot of October 21, 2026^65^, prioritizing approved symbols and uniquely resolved aliases or curated mappings. Ambiguous aliases were excluded from gene-set memberships without reducing the fixed universe.

For GO Molecular Function, 1,918 MSigDB 2026.2.Hs gene sets^43–45^ were tested separately for each Gate group against the remainder of the fixed universe using one-sided Fisher exact tests for enrichment. Benjamini–Hochberg correction was applied across all 1,918 tests within each group, including 107 sets with no background members assigned P = 1.

For Hallmark ORA, all 50 MSigDB 2026.2.Hs Hallmark sets^32^ were tested for enrichment in Gate #2 using one-sided Fisher exact tests, with Benjamini–Hochberg correction across the 50 tests. Conditional odds ratios were reported.

TRRUST v2 human regulons^34^ were included if they contained at least 10 target genes in the fixed background, yielding 102 regulons. One-sided Fisher enrichment tests were performed, with Benjamini–Hochberg correction across a combined family of 24,510 tests: 24,403 MSigDB 2026.1.Hs sets, five additional gene sets, and 102 TRRUST regulons. The TRRUST, IEG, and DPRG results used this common correction family.

IEG and DPRG sets contained 45 and 50 unique genes after mapping the 49 and 58 genes reported by Tullai et al.^33^ to Ensembl gene annotations.

### Promoter-proximal Pol II enrichment

Promoter-proximal Pol II enrichment before serum restimulation was analyzed using Guo MCF10A PRO-seq (GSM4914401) and total Pol II ChIP-seq (GSM4915953)^36^; Hyder Pol II (RPB3) ChIP-seq (GSM8253808/GSM8253809) and PRO-seq (GSM8253859/GSM8253860; processed GSE266692 track) from serum-starved, DMSO-treated KAP1-dTAG knock-in HCT116 cells (clone D5)^37^; and total Pol II ChIP-seq from serum-starved, DMSO-treated HCT116 cells (GSM1545657/GSM1545658; GSE63314)^66^. Guo tracks were single profiles, whereas Hyder profiles were author-merged from two registered replicates.

Promoter-proximal enrichment was calculated as the log2 ratio of mean coverage from TSS −50 to +300 bp to mean coverage from TSS +500 bp to TES −500 bp, oriented relative to transcription. Annotation used GENCODE v50 and MANE v1.5^67,68^. Genes were required to have a transcript span ≥2 kb, no promoter or gene-body overlap with blacklist regions^69^, positive promoter and body coverage, at least 100 positive body bases, and body signal at or above the 20th percentile of positive background body signal among structurally eligible genes. Ratios were calculated from strand-merged or unsigned processed coverage without pseudocounts or ChIP/input correction.

Gate #2 and non-Gate #2 gene distributions were compared using two-sided Mann–Whitney tests with asymptotic inference, tie handling, and continuity correction. Benjamini–Hochberg correction covered all five pre-restimulation comparisons, including the undisplayed Chen et al. Pol II ChIP-seq comparison^66^. The four displayed comparisons contained 1,015/6,515 Gate #2/non-Gate #2 genes for Guo PRO-seq, 1,001/6,448 for Guo ChIP-seq, 945/6,612 for Hyder PRO-seq, and 943/6,651 for Hyder ChIP-seq.

### Public stimulus-response time courses

Five public stimulus-response time courses were analyzed as follows.

### GSE6783 HeLa EGF

HeLa cells were treated with EGF (10 ng/mL) after 24 h of serum deprivation, with samples collected at 0, 20, 40, 60, 120, 240, and 480 min^38^. One array was available per time point. GEO-deposited MAS5 intensities were used; for each gene, the probe with the highest baseline intensity among baseline-present probes was selected. Responses were calculated relative to 0 min.

### GSE9120 keratinocyte IL-1α

Keratinocytes were treated with IL-1α (25 ng/mL) after 24 h in KBM and sampled at 1, 4, 24, and 48 h, with one treated and one time-matched untreated array per time point^39^. The eight CEL files were jointly processed by RMA^70^, and responses were calculated as log2-intensity differences from time-matched controls.

### GSE202436 HBEC TNF-α

HBECs from five donors (K, P, S, V, and T) were treated with TNF-α (10 ng/mL) for 1, 2, 6, or 18 h^41^. GEO-deposited kallisto gene TPM values were used to calculate donor-paired log2 ratios relative to time-matched NS vehicle controls, which were then averaged across donors.

### GSE112372 macrophage LPS

Macrophages from four donors (1, 2, 10, and 11) were treated with LPS (100 ng/mL) and sampled at 0, 0.5, 3, 8, and 16 h; donor 10 was absent at 16 h^42^. GEO-deposited RSEM expected counts were analyzed using edgeR TMM normalization and a GLM with donor and time terms^63,64,71^, with responses calculated relative to untreated 0 h.

### GSE17708 A549 TGF-β1

A549 cells were treated with TGF-β1 (5 ng/mL) after 24 h of serum starvation and sampled at 0, 0.5, 1, 2, 4, 8, 16, 24, and 72 h^40^. Three experiments were available, except at 2 h, which had two. The 26 CEL files were jointly processed by RMA^70^, and paired experiment-level log2 differences from 0 h were averaged. The 72-h point was omitted from the main figure but retained in the multiple-testing correction family.

### Time-course enrichment and cross-series responses

At each time point, evaluable genes in the union of Gate #1, Gate #2, and Others were ranked by signed log2 fold change in descending order. The top-five-percent cutoff was rank ceiling(0.05N), including all ties at the boundary, without an additional positivity filter. The denominator for each proportion was the corresponding evaluable Gate group, and Wilson 95% binomial confidence intervals were calculated over genes. Groups were compared using two-sided Fisher exact tests. Within each dataset, Benjamini–Hochberg correction covered the analyzed conditions, all time points, and the FC≥1.5, top-5%, and top-10% rules.

For Figure 6F, a gene was classified as responding in a series if it ranked in the top 5% at any sampled time within 0 < t ≤4 h. This analysis used 353 Gate #2 and 3,736 non-Gate #2 genes evaluable across all five series.

## Data Availability

RNA-sequencing and ribosome-profiling data generated in this study have been deposited in the Gene Expression Omnibus (GEO) under accession number GSE354450. Mass spectrometry proteomics data generated in this study have been deposited with the ProteomeXchange Consortium via the jPOST repository^72^ under accession numbers PXD087119 and JPST005072.

Any additional information required to reanalyze the data reported in this paper is available from the corresponding author, Hirokazu Nakatsumi, upon request.

## Materials Availability

Requests for resources and reagents should be directed to the corresponding author, Hirokazu Nakatsumi. Cell lines and plasmids generated in this study are available upon request and may require a completed materials transfer agreement with Nagoya City University.

## Acknowledgments

We thank the Research Equipment Sharing Center at Nagoya City University for providing access to shared research equipment, the members of the Tokai TOR Conference (ToToCo) for helpful discussions, and all members of the Shirane laboratory for support. This work was supported by JSPS KAKENHI Grant Numbers JP21K07155 and JP26K09319 (to H.N.) and JP20H03255, JP20H04907, and JP24K02122 (to M.S.); and by the Takeda Science Foundation, the Suzuken Memorial Foundation, the Toyoaki Scholarship Foundation, the Research Foundation for Pharmaceutical Sciences, and the Ichihara International Scholarship Foundation (to H.N.).

## Author Contributions

Conceptualization, H.N.; Methodology, H.N., K.O., and Y.H.; Investigation, H.N., K.I., T.M. A.H., T.A., Y.I., A.M., M.M., and K.I.N.; Formal analysis, H.N.; Data curation, H.N.; Visualization, H.N.; Writing – original draft, H.N.; Writing – review & editing, all authors; Supervision, K.O., Y.H., and M.S.; Funding acquisition, H.N. and M.S.

## Declaration of Generative AI Use

During the preparation of this work the authors used Claude (Anthropic), ChatGPT (OpenAI), and Gemini (Google) in order to improve the clarity, concision, and readability of the manuscript text. After using these tools, the authors reviewed and edited the content as needed and take full responsibility for the content of the publication.

## Supplementary Information

**Figure S1.**
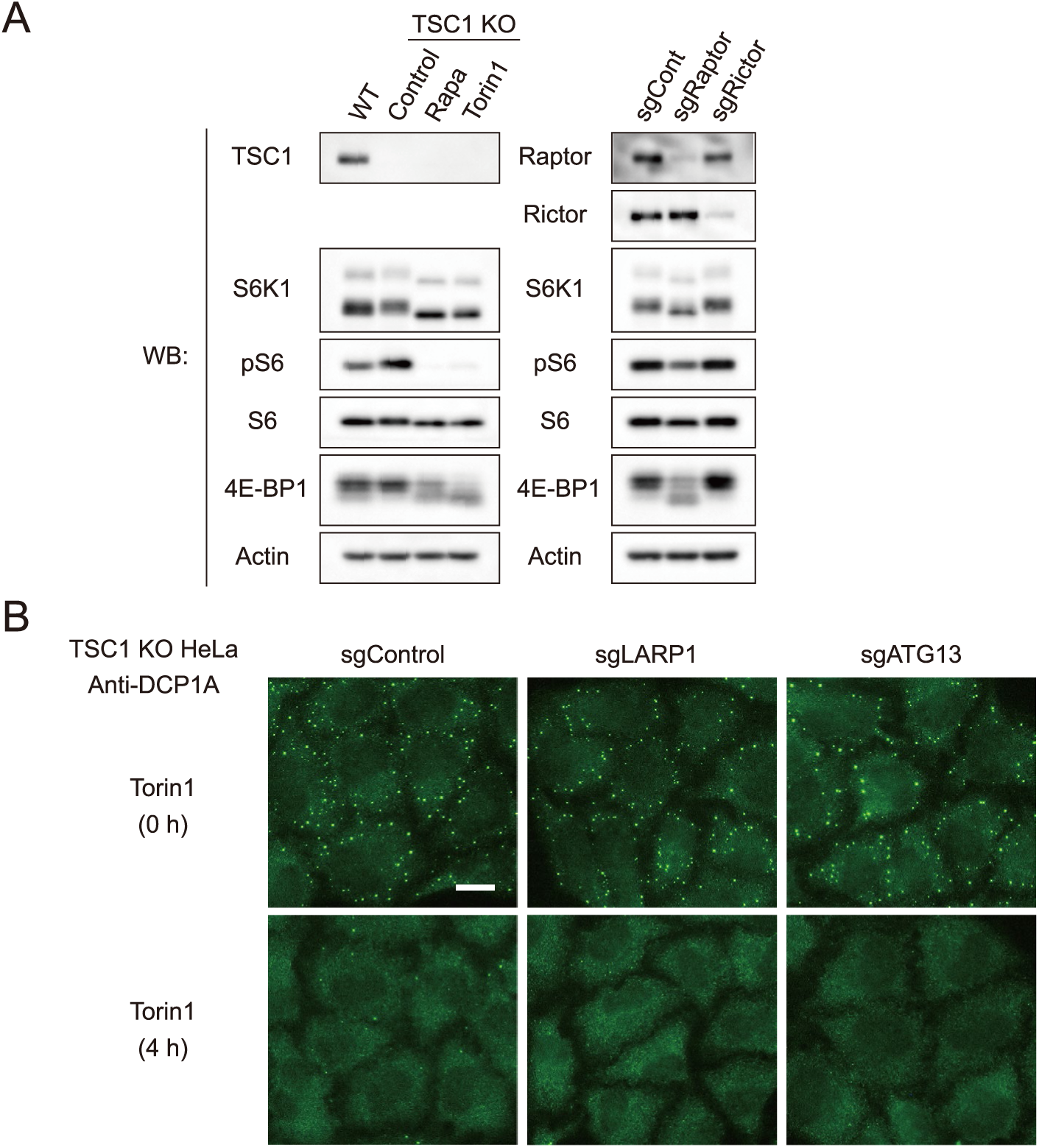
Validation of mTORC1 perturbations and P-body responses, related to Figure 1. (A) Immunoblotting for TSC1, Raptor, Rictor, S6K1, phosphorylated S6 (pS6), S6, 4E-BP1, and actin in the cells used in Figures 1B and 1C, confirming target depletion and the expected changes in mTORC1 signaling. (B) DCP1A immunofluorescence in TSC1 KO HeLa cells transduced with sgControl, sgLARP1, or sgATG13, before and 4 h after Torin1 treatment (100 nM). Scale bar, 20 µm.

**Figure S2.**
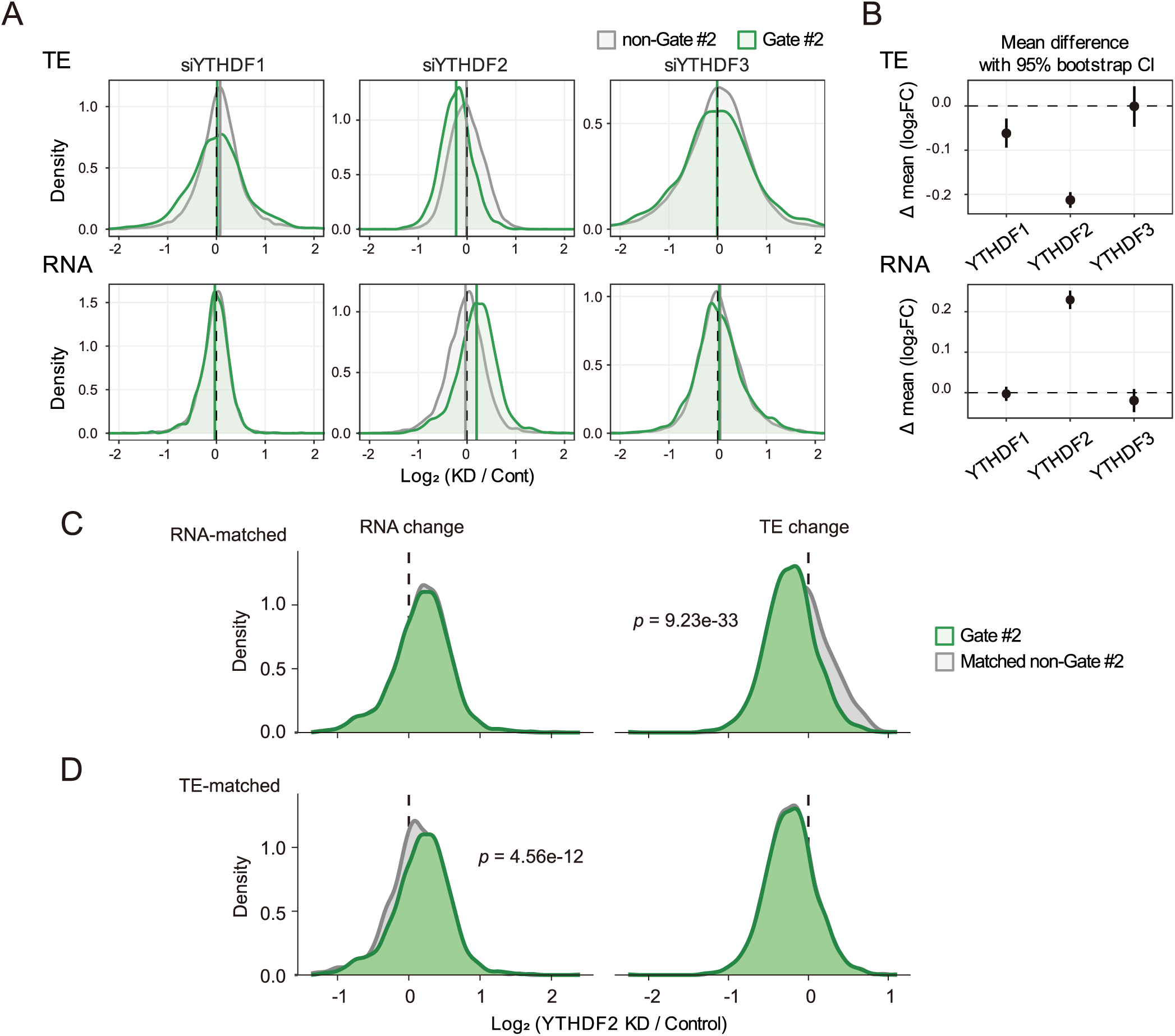
Changes in TE and RNA abundance after YTHDF knockdown and matched-control analyses, related to Figure 5. (A) Distributions of TE (top) and RNA abundance (bottom) log2 fold changes (knockdown/control) in Gate #2 versus non-Gate #2 for each YTHDF knockdown^14,29,30^. (B) Differences in mean log2FC between Gate #2 and non-Gate #2 for each knockdown. Error bars, percentile 95% bootstrap CIs. (C) For YTHDF2 knockdown, a non-Gate #2 control set matched to Gate #2 by RNA abundance log2FC; the matched sets are compared for RNA log2FC (left) and TE log2FC (right). (D) As in (C), with the control set matched by TE log2FC; the matched sets are compared for TE log2FC (right) and RNA log2FC (left). The primary comparisons are TE changes after RNA matching (C, right) and RNA changes after TE matching (D, left); the other panels show matching quality. Two-sided paired Wilcoxon signed-rank tests on matched gene pairs, unadjusted P values.

**Table S1.**
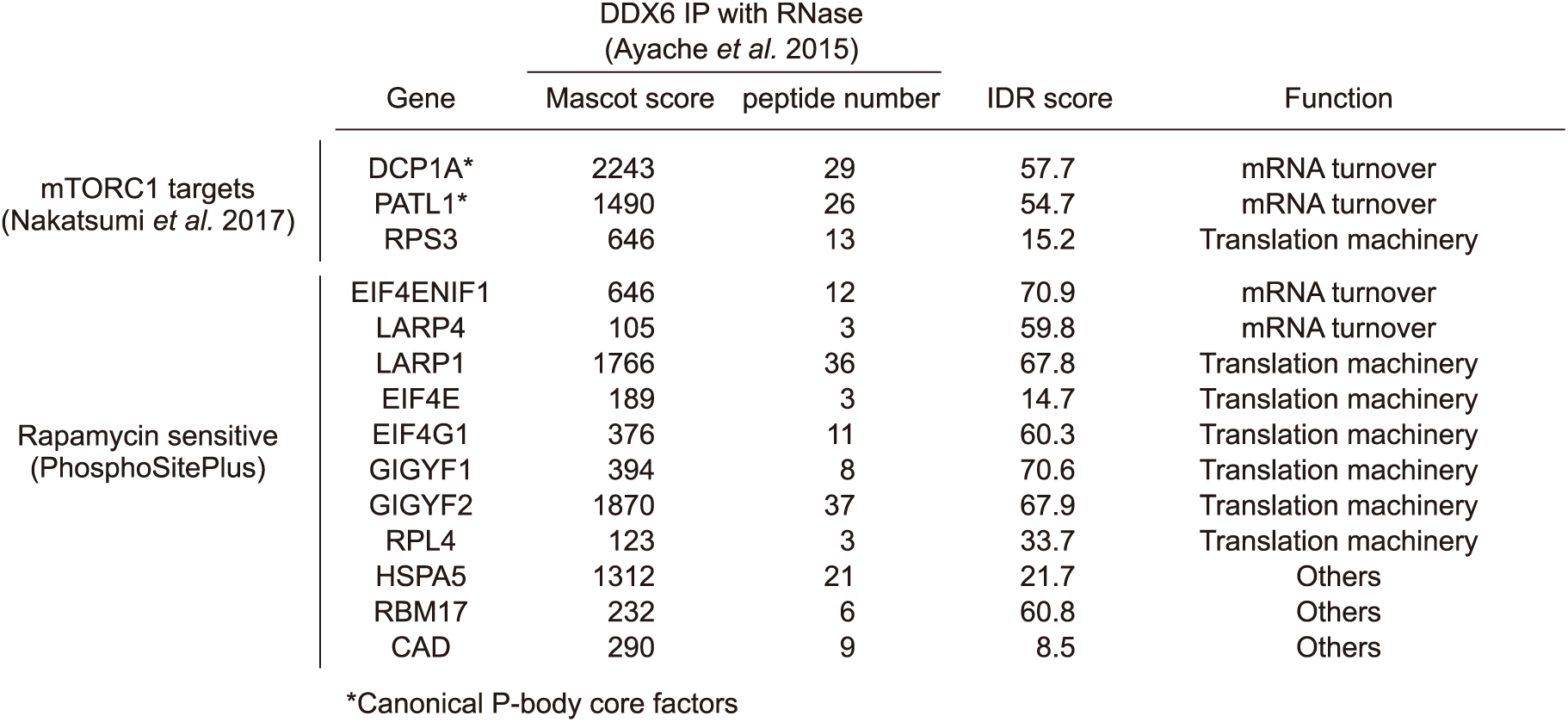
DDX6-interacting proteins overlapping mTORC1-associated or rapamycin-sensitive phosphorylation sets, related to Figure 2. Fourteen DDX6-interacting proteins identified by RNase-treated DDX6 immunoprecipitation^21^ that overlap either mTORC1-associated phosphorylation targets^20^ or rapamycin-sensitive phosphoproteins from PhosphoSitePlus^22^. Mascot scores, numbers of identified peptides, intrinsic-disorder scores, and functional categories are shown. Asterisks indicate canonical P-body core factors.

|  | Gene | DDX6 IP with RNase<br>(Ayache *et al.* 2015) |  | IDR score | Function |
|---|---|---|---|---|---|
|  |  | Mascot score | peptide number |  |  |
| mTORC1 targets<br>(Nakatsumi *et al.* 2017) | DCP1A* | 2243 | 29 | 57.7 | mRNA turnover |
|  | PATL1* | 1490 | 26 | 54.7 | mRNA turnover |
|  | RPS3 | 646 | 13 | 15.2 | Translation machinery |
| Rapamycin sensitive<br>(PhosphoSitePlus) | EIF4ENIF1 | 646 | 12 | 70.9 | mRNA turnover |
|  | LARP4 | 105 | 3 | 59.8 | mRNA turnover |
|  | LARP1 | 1766 | 36 | 67.8 | Translation machinery |
|  | EIF4E | 189 | 3 | 14.7 | Translation machinery |
|  | EIF4G1 | 376 | 11 | 60.3 | Translation machinery |
|  | GIGYF1 | 394 | 8 | 70.6 | Translation machinery |
|  | GIGYF2 | 1870 | 37 | 67.9 | Translation machinery |
|  | RPL4 | 123 | 3 | 33.7 | Translation machinery |
|  | HSPA5 | 1312 | 21 | 21.7 | Others |
|  | RBM17 | 232 | 6 | 60.8 | Others |
|  | CAD | 290 | 9 | 8.5 | Others |

\*Canonical P-body core factors

**Table S2.** P-body localization and CDS-m⁶A enrichment scores and Gate assignments, related to Figure 3. The table lists Ensembl gene IDs, gene symbols, P-body localization and CDS-m⁶A enrichment scores (log2 fold changes), their corresponding Z scores, and a combined score calculated as the sum of the two Z scores for 11,057 genes. Gate assignments were determined using the thresholds described in Materials and Methods.

**Table S3.** Reagents and resources used in this study.

